# Elsewhere or Blanked? Ongoing mental states are regulated by pupil-linked arousal and attentional style in healthy ageing

**DOI:** 10.1101/2022.07.08.499379

**Authors:** Matthieu Koroma, Aurèle Robert de Beauchamp, Sepehr Mortaheb, Paradeisios Alexandros Boulakis, Christine Bastin, Athena Demertzi

## Abstract

Spontaneous thinking significantly relies on attention and arousal. As these cognitive faculties change with age, we aimed at providing a comprehensive account to ongoing mental states in seniors, testing how these are influenced by attentional control and arousal. Using experience sampling at rest, 20 senior (65-75yrs) and 20 young participants (20-30yrs) were prompted to report mind-wandering (MW), sensory-related thoughts (S), and the newly introduced state of mind blanking (MB). Attentional control was assessed with the Attentional Style Questionnaire, and arousal with continuous monitoring of pupil diameter. Both age groups showed equally high occurrences of MW compared to MB or S. For young responders, we replicated that MW was more prevalent in easily-distracted participants and that it associated with higher arousal. In seniors, though, MB was more prevalent in easily-distracted participants, and it was associated with higher arousal, reversing the pattern found in young adults and focused seniors. Overall, our results show that attentional control and arousal jointly regulate ongoing mental states in an age-dependent manner and uncover the presence a specific profile of ongoing mental state regulation in healthy aging, being a potentially critical marker of age-associated diseases.

## Introduction

Healthy aging has been widely described in terms of variant modifications in the cognitive and behavioral domain (Craik & Salthouse, 2008; Murman, 2015), and recently it started being characterized in terms of ongoing thinking as well (Jackson & Balota, 2012; Seli et al., 2017). Ongoing thinking refers to the ability to entertain and transition among various mental states, which are described in terms of content (what the state is “about”) and the relation we bear to this content, such as imagining, remembering, and fearing (Christoff et al., 2016). As ongoing thoughts, especially those that derail from the task at hand, can occupy 10%-60% of waking life (Killingsworth & Gilbert, 2010; Seli et al., 2018), the relationship between cognitive changes and ongoing mental states may provide useful insights not only for typical ageing but also for age-associated diseases (Gyurkovics et al., 2018) .

During spontaneous thinking, the way we transition among mental states is heavily based on how we allocate our attentional resources for cognitive control. An indicative example is mind-wandering (MW), a mental state which consists in diverted attention from performing a task to entertain inner thoughts, fantasies, feelings, and other musings (Smallwood & Schooler, 2006), either spontaneously or deliberately (Seli et al., 2017). It was previously shown that, as attention disengages from perception (Schooler et al., 2011), MW can happen with awareness (“tune out”) or without awareness (“zone out”) (Smallwood et al., 2007). Or it can take the form of an active MW state, during which attention is focused toward an internal event, or of an off-focus state, during which there is no specific thought and the attentional focus is broadened (Mittner et al., 2016).

This latter model predicts that the off-focus state should be characterized by higher tonic (baseline) pupillary diameter to account for this explorative mode to allow the selection of a new thought. Recently, the relation between the tonic pupillary diameter and MW by taking the different states of attention into account was examined with a sustained-attention-to-response task (Jubera-García et al., 2020): by considering thoughts with high intensity (clear focus) and with low intensity (no clear focus), the results on tonic pupillary diameter were not in line with the model’s predictions and failed to replicate between the experiments. At the same time, behavioral responses were related to the intensity of thoughts, with behavioral variability being smallest when participants were having on-task thoughts, intermediate for “off-focus” thoughts, and highest for active MW. To interpret these findings, the authors relied on the pupil results and suggested that the off-focus state might be more complex than it was initially considered, such that it does not only refer to the intensity of thoughts, but also to moments of no content (Jubera-García et al., 2020).

Such off-moments with no particular content have been recently quantified, as the experience that the mind is absent or went “blanked” (Kawagoe et al., 2019; Ward & Wegner, 2013). Mind blanking (MB) has been quantified as a mental state happening by default only recently, as it was reported with low but equal probabilities along the total acquisition time (Mortaheb et al., 2022). Operationally, MB has been defined as “reports of reduced awareness and a temporary absence of thought (empty mind) or lack of memory for immediately past thoughts [which] can be considered as the phenomenological dimension of a distinct kind of attentional lapse” (Andrillon et al., 2019, p2). Such attentional lapses linked to MB were shown to correlate with the presence of transient slow wave activity indicative of low cortical arousal even during wakefulness (Andrillon et al., 2021). In the larger scale of the fMRI signal, the arousal hypothesis was also put forward as the authors identified that MB reports were linked to a configuration pattern characterized by high global signal amplitude (Mortaheb et al., 2022), a proxy for decreased arousal (Liu et al., 2015, 2018; Wong et al., 2013).

Taken together, studies in young participants indicate that attention and arousal are important cognitive faculties when investigating ongoing mentation. With ageing, these faculties have been shown to change. Selective attention changes with age were shown by increments in target identification time and error rate (Plude & Hoyer, 1986). Differences in divided attention (Wright, 1981) have also been reported in terms of simple immediate attention span, selectivity, capacity to inhibit interference of non-pertinent signals, and attentive shifting (Commodari & Guarnera, 2008). Additionally, aging affects vigilance in sustained attention tasks (Berardi et al., 2001) and is associated with decreased pupil size both in light and dark conditions (Birren et al., 1950). Collectively, attentional resources and arousal changes are concomitant in senior individuals and influence task performance (Craik & Salthouse, 2008).

In terms of ongoing mentation, an age-related reduction in the frequency of MW has been reported (Gyurkovics et al., 2018; Jackson & Balota, 2012; Seli et al., 2017). MB reports, though, have not been characterized. For a more comprehensive characterization of ongoing thinking in healthy seniors, we evaluated the behavioral and physiological profiles of mental states during rest and checked their potential regulation by age-dependent attentional and arousal modifications. Based on the aforementioned literature, we hypothesized that: 1) MW will be associated with lower attentional control, as indexed by a higher score on the Attention Style Questionnaire; 2) MB will be associated with lower arousal, as indexed by a smaller pupil size; 3) Seniors will show lower rates of MW and higher rates of MB as compared to younger participants due to overall higher task focus (Maillet et al., 2020; Seli et al., 2021) and smaller pupil-linked arousal (Birren et al., 1950).

## Methods

### Participants

Forty-two French-speaking adults were recruited by means of advertisement in University of Liège and University of the Third Age. Participants were split in two groups: *seniors* (65-75yrs) and *young* participants (20-30yrs). One senior and one young participant were excluded due to technical problems with ocular recordings, resulting in a final cohort of 20 seniors (14 females, mean age=69.3±3.1) and 20 young participants (15 females, mean age=25.4±2.4). This sample size allows for the detection of small effect sizes (f=0.1) with a power of 0.8 and an alpha of 0.05 for a within-between repeated measures design with two groups, forty measures and a correlation among repeated measures of 0.33 in a decision among three options. All individuals had normal or corrected-to-normal vision and audition with no history of psychiatric or neurological disorders. The study was approved by the ethics committee of the Faculty of Psychology, Speech Therapy and Educational Science of the University of Liège. All participants provided written informed consent prior to the study.

### Procedure

Participants were invited to sit comfortably in a dimly lit room in front of a computer screen and a keyboard. They were instructed to use earphones (ER-3C, Etymotic Research) to ensure auditory isolation from external sounds and reduction of auditory distraction from potential surrounding sound sources. Participants wore oculometric glasses (Phasya Drowsimeter R100) which allowed for pupil size monitoring, which was considered as an objective physiological marker of arousal (Alnaes et al., 2014). Before the experiment, participants were asked to fixate a point on the screen for a few seconds to calibrate the eye recording system.

Participants were explained the aim of the task and were given the definitions of “Mind-Wandering” (MW), “Mind-Blanking” (MB), and “Sensations” (S). MW was defined as all types of thoughts, related or not to the present environment, such as “I think about the screen I am looking at” or “I am thinking about what I am going to eat tonight”. MB was defined as “an empty mind”, “thinking about nothing” but also “no memory of mental content”. S was defined as the sensorial acknowledgment related to the five senses of one or more stimuli without any thought attached to it, such as “I heard someone’s voice without thinking”. The definitions were repeated until they were correctly distinguished and understood.

The experiment was designed and presented using the PsychoPy toolbox (PsychoPy3 v.2020.2.10). The task utilized the probe-caught experience-sampling method (Smallwood & Schooler, 2006) consisting of the participant instructed to maintain their gaze on the screen while their mind was allowed to go freely and 40 auditory prompts (1s tone-pip, 440 Hz frequency, played at 50 dB, sampled at 44000 Hz) presented at random time intervals, ranging between 30 to 60 sec. The prompts invited participants to report on their mental state as it was prior to the tone by using one out of three options: “MW”, “MB”, and “S” (Figure 1).

**Figure 1.**
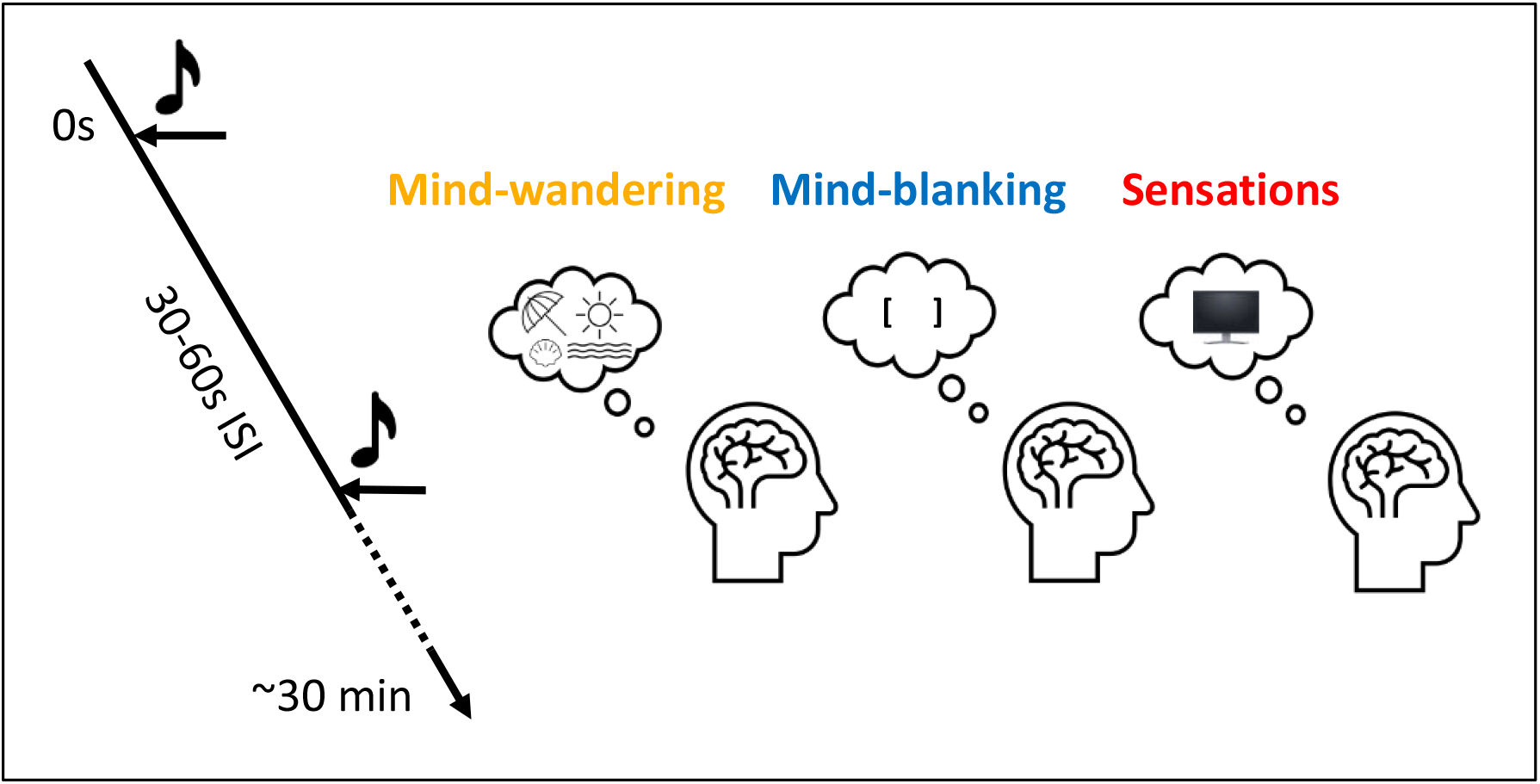
**Experience-sampling task**. Participants’ ongoing resting state was interrupted by 40 auditory prompts randomly appearing with an inter-stimulus interval (ISI) 30 to 60 sec. The prompt invited them to report their mental state as it was beforehand by choosing among “Mind-Wandering”, “Mind-Blanking”, and “Sensations” (*e.g.,* the view of the experiment screen).

The experiment started upon the participant’s keypress. The computer screen was maintained grey throughout the experiment except after the sound probes when the response options and their corresponding key press appeared on the screen. After the response was provided, the screen turned grey again. Participants were instructed to keep their gaze on the screen and their eyes open during the whole experiment. The total duration of the experiment was about 30 minutes.

### Questionnaires

Upon task completion, participants were administered the Attentional Style Questionnaire (ASQ), the validated French version Generalized Anxiety Disorder – 7 (GAD-7), and the Mini Mental State Examination (MMSE, only for seniors). The ASQ is a self-assessment questionnaire measuring the capacity of an individual to maintain attention on task-related stimuli and not to be distracted by interfering stimuli (Smallwood & Schooler, 2006). A lower ASQ score reflects a more focused attentional style. Only psychometrically validated items of the questionnaire were used (Kraft et al., 2020), forming 12 items ranked on a Likert scale ranging from “strongly disagree” to “strongly agree”.

We also considered anxiety symptoms as a potential confound of our results as they have been reported to affect attentional abilities, pupil size variations and the occurrence of ongoing mental states (Makovac et al., 2019). Participants filled the GAD-7, a 7-item self-rated questionnaire designed to assess the intensity of anxiety symptoms (Spitzer et al., 2006). Each item is rated on a Likert scale from 0 (not at all) to 4 (nearly every day).

The Mini Mental State Examination (MMSE; Folstein et al., 1975) quantified general cognitive functions and check the absence of any major neurocognitive disorder. The MMSE consists of 6 categories (orientation, learning, attention and calculation, reminder, language, and constructive praxis). Although the classification of the obtained score depends on the person’s socio-cultural level, a score of 24 or less on a maximum of 30 indicates mild dementia.

### Behavioral data analysis

Collected behavioral data concerned mental state reports and reaction times. Prevalence of mental state was determined in terms of overall occurrence rate for each mental state (MW, MB, S), reaction times with regards to button presses, the distribution of appearance of mental state reports across the experiment, and inter-state transition dynamics.

To test whether mental state occurrence rate depended on the type of mental state, age group and attentional style of anxiety, we used a generalized linear-mixed model with a binomial distribution, the number of trials (n=40) as weighting factor and participants defined as random effect factors. The influence on reaction times of mental states, age group and attentional style or anxiety was assessed with a generalized linear mixed-effect model with a gamma distribution and an inverse link function, with participants defined as random effect factors. In both cases, the glmer function in R was used.

Post-hoc paired non-parametric Wilcoxon signed-rank tests with false discovery rate (FDR) correction for multiple comparisons were used to assess the difference in occurrence rates across mental states within groups. Non-parametric unpaired Wilcoxon rank-sum tests with FDR correction for multiple comparisons were used to assess the difference in the rates of occurrence of mental states, attentional style and anxiety between groups. Effect sizes were computed using the formula: r= Z/√n, where Z is the Z-value and n the number of participants (Pallant, 2020). The correlation between ASQ and GAD scores was assessed with Pearson’s correlation coefficient (R).

Distribution of ongoing mental states across the entire experiment was statistically assessed by comparing it to a uniform distribution using Chi-squared test. Inter-state transition dynamics was assessed by computing the probability of transition between mental states for each participant and testing for differences across age groups using Wilcoxon-rank sum tests with FDR correction for multiple comparisons.

### Physiological analysis

Collected physiological data concerned pupil dilation, eye-lid gap, and eye-movements recorded at 120Hz using the Drowsilogic v 4.3.9. software for each mental state (MW, MB, S).

Time series of physiological signals were extracted over the last 30 seconds preceding probe onset for each trial. Missing data due to eye blinks were linearly interpolated using the most neighboring values. Differences between mental states for each age group were compared using the non-parametric cluster-level permutation test function (alpha-level=0.05, n=1023 permutations) as implemented in the MNE package (version 0.23.0, https://mne.tools/).

The interactions between mental states, age and attentional style on physiological signals over the last 30 seconds of each trial were assessed with linear mixed effect modelling using the lmer function in R and participant defined as random effect. Post-hoc investigations of the interaction between attentional resources and arousal across mental states and age, physiological signals were investigated after averaging signals over the last 30 seconds and normalizing z-scoring values for each participant. Post-hoc paired non-parametric Wilcoxon signed-rank tests with false discovery rate (FDR) correction for multiple comparisons were used to assess the difference in occurrence rates across mental states within groups. Correlations between ASQ scores and physiological variables across participants was assessed for each mental state and age group with Pearson’s correlation tests and corrected for multiple comparisons using the false discovery rate (FDR).

## Results

### Ongoing mental state profiles

All senior participants were free of cognitive decline (MMSE mean score= 29.3±0.8, range= 28-30). A higher occurrence rate of MW over MB and S was found in both groups (MW vs. MB: z(119,108)=8.14, p<0.001, MW vs. S: z(119,108)=6.12, p<0.001). Post-hoc tests confirmed the higher occurrence of MW over MB and S in senior (Wilcoxon signed-rank test, MW vs. MB: r=0.60, p<0.001, MW vs. S: r=0.62, p<0.001) and in young adults (MW vs. MB: r=0.59, p<0.001, MW vs. S: r=0.62, p<0.001) (22).

A similar MW predominance was revealed with regards to reaction times. MW was reported faster than MB (z(108,40)=-2.78, p=0.005) and S (z(108,40)=-2.09, p=0.037), while MB and S were reported equally fast (z(108,40)=-2.77, p=0.622) (Figure 2, upper panel). RTs in seniors and young participants did not differ with regards to mental state (seniors vs. young with MW vs. MB: z(108,40)=1.63, p=0.101, MW vs. S: z(108,40)=0.85, p=0.394, MB vs. S: z(108,40)=-0.67, p=0.502), suggesting that mental states were categorized with similar efficiency across age groups (Figure 2, lower panel).

**Figure 2.**
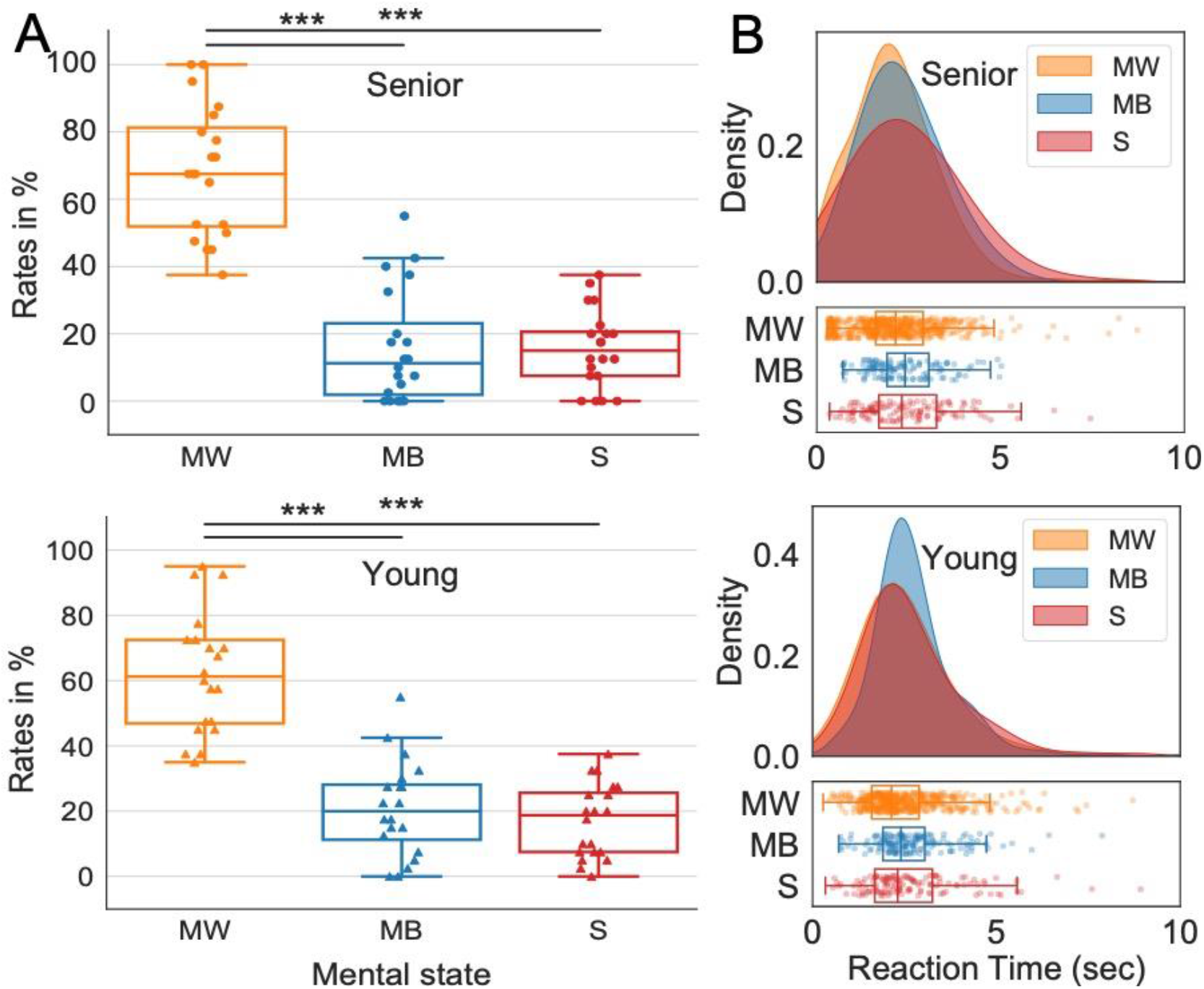
Rates of ongoing mental states in senior and young participants. (A) Mind-wandering (MW) predominates over Mind-Blanking (MB) and Sensations (S) in all participants independently of age group. (**B)** Distribution of reaction times for mental states for senior (upper panel) and young participants (lower panel) shows that MW was reported faster than other mental states, with similar efficiency across age groups. *Notes*: Boxplots: mean (horizontal line), first and third quartile (box), as well as first and ninth deciles (vertical line) for Mind-Wandering (MW, yellow), Mind-Blanking (MB, blue), and Sensations (S, red) represented for senior (solid) and young participants (dotted). Individual data points of each are represented as circles (senior) and triangles (young), ***p<0.001 Wilcoxon rank-sum tests. Distribution of reaction times modelled with a continuous probability density curve (kernel density) for each mental state.

Finally, we checked whether mental states were reported differently across time between age groups. By investigating the distribution of mental states over the experiment, no evidence for unequal distribution of response types over time for all mental states in each age group was found (Chi-squared tests, p> 0.05) (Figure S1). Probabilities of transition between mental states were also similar across both age groups (Wilcoxon rank-sum tests, p> 0.05) (Figure S2), providing no evidence for changes in the inter-state dynamics across age groups.

### Attentional style

Seniors self-reported lower scores on the Attentional Style Questionnaire (ASQ) than younger participants (37.2±[33.7, 40.7] vs. 46.6±[43.5, 49.7], r=- 0.56, p<0.001, Figure 3), indicating a higher focus ability for senior participants. The interaction between attentional style, rates of ongoing mental state, and age group was identified between MW and MB reports (z(108,40)=-6.15, p<0.001), MW and S reports (z(108,40)=5.58, p<0.001) but not between MB and S (z(108,40)=0.381, p=0.703). Therefore, we investigated the association between the ASQ score and the occurrence of ongoing mental states across age group.

**Figure 3.**
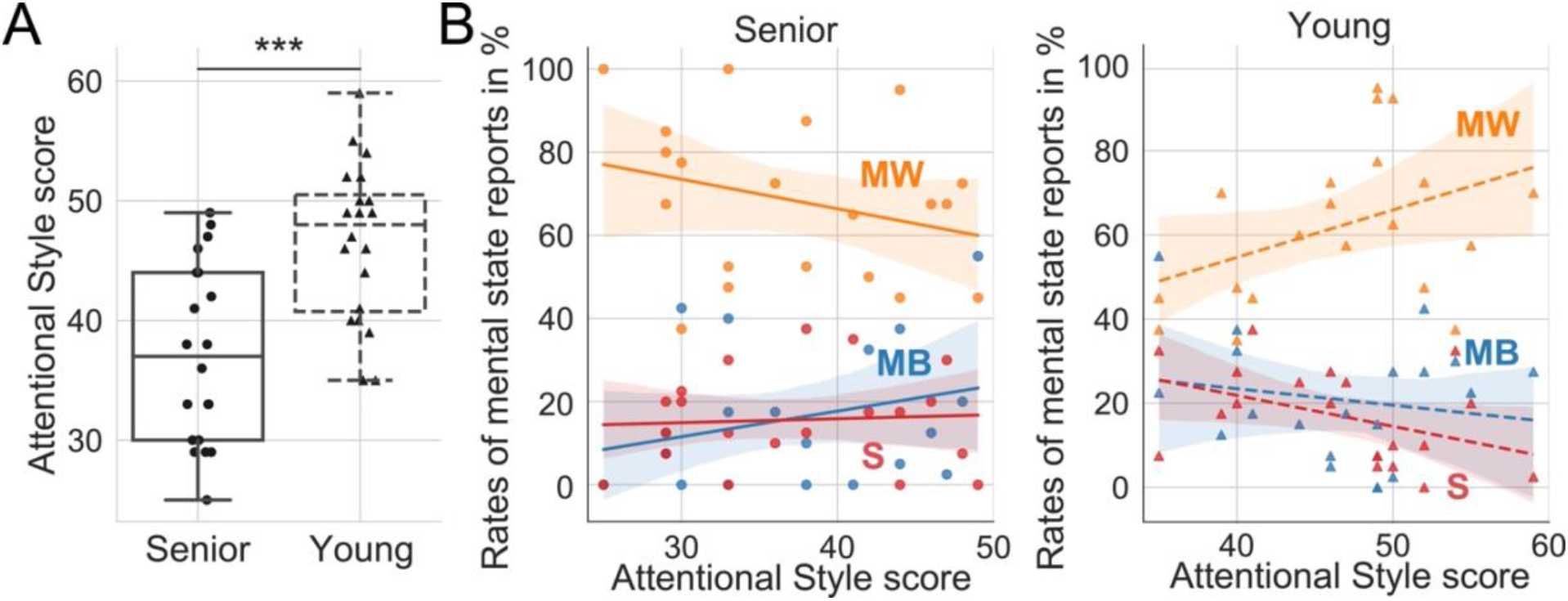
Age-related attentional style reverses the regulation of ongoing mental states in seniors. (A) Seniors self-reported lower scores on the Attentional Style Questionnaire (ASQ) than younger participants, indicative of higher attentional focus. (**B**) In seniors, lower ASQ scores were associated with higher MW (left panel) and the reverse pattern was found in younger participants who showed that lower ASQ scores were linked to lower MW (right panel), suggesting that age reverses the relationship between attentional focus and rates of ongoing mental states. *Notes*: Boxplots: mean (horizontal line) and first and third quartile (box), as well as first and ninth deciles (vertical line) of the Attentional Style Questionnaire (ASQ) score are represented (solid: aged, dotted: young). Individual data points are represented in circles: (senior) and triangles (young). Significant differences across conditions are reported as solid black lines (Wilcoxon rank-sum test, ***p<0.001). Correlation plots: The lines (solid: senior, young: dotted) represent the correlation between the rate of Mind-Wandering (MW): yellow, Mind-Blanking (MB): blue, Sensations (S): red, and the ASQ score with shaded area as confidence intervals for both senior (left panel) and young (right panel) participants. Individual data points of each are represented as circles (senior) and triangles (young).

In seniors, lower ASQ scores were linked to higher MW over MB (z=4.65, p<0.001) and higher MW over S rates (z=2.38, p<0.01) (Figure 3, left panel). The reverse pattern was found in younger participants who showed that lower ASQ scores were linked to lower MW over MB (z=-4.07, p<0.001) and S rates (z=-5.38, p<0.001) (Figure 3, right panel), meaning that age reverses the relationship between attentional control and ongoing mental states.

We further checked whether our results based on attentional style could be confounded by the effect of anxiety on the occurrence of mental states. We first confirmed that lower attentional control (high ASQ scores) correlated with higher anxiety scores (Pearson’s correlation test: R(37)=0.33, p=0.043, Figure S3). We also found that seniors had a lower anxiety score than young participants (4.58±[2.45, 6.71] vs. 7.25±[4.98, 9.52], r=-0.31, p=0.025) (Figure S4A); however, anxiety scores did not predict significantly rates of mental states nor interacted with the type of mental state and age groups (mixed-model effects: p- values > 0.05) (Figure S4B). This excludes anxiety as a potential confound explaining the influence of attentional style on the occurrence of ongoing mental states.

### Arousal

Across the experiment, seniors had smaller pupil size when reporting MW than MB ([-30 to 0]s, F=7.93, p=0.004), and MW than S ([-30 to -15.8s]s : F=10.67, p=0.006; [-15.6 to 0]s: F=7.39, p=0.009), while no time cluster were found for young participants with regards to pupil size variations across mental state (Figure 4). Linear mixed-effects models revealed a significant interaction between age and pupil size difference between MW and MB (t(108,40)=-2.93, p=0.005), as well as between MW and S (t(108,40)=-3.57, p<0.001). These results confirm that physiological signatures of ongoing mental states differ with age.

**Figure 4.**
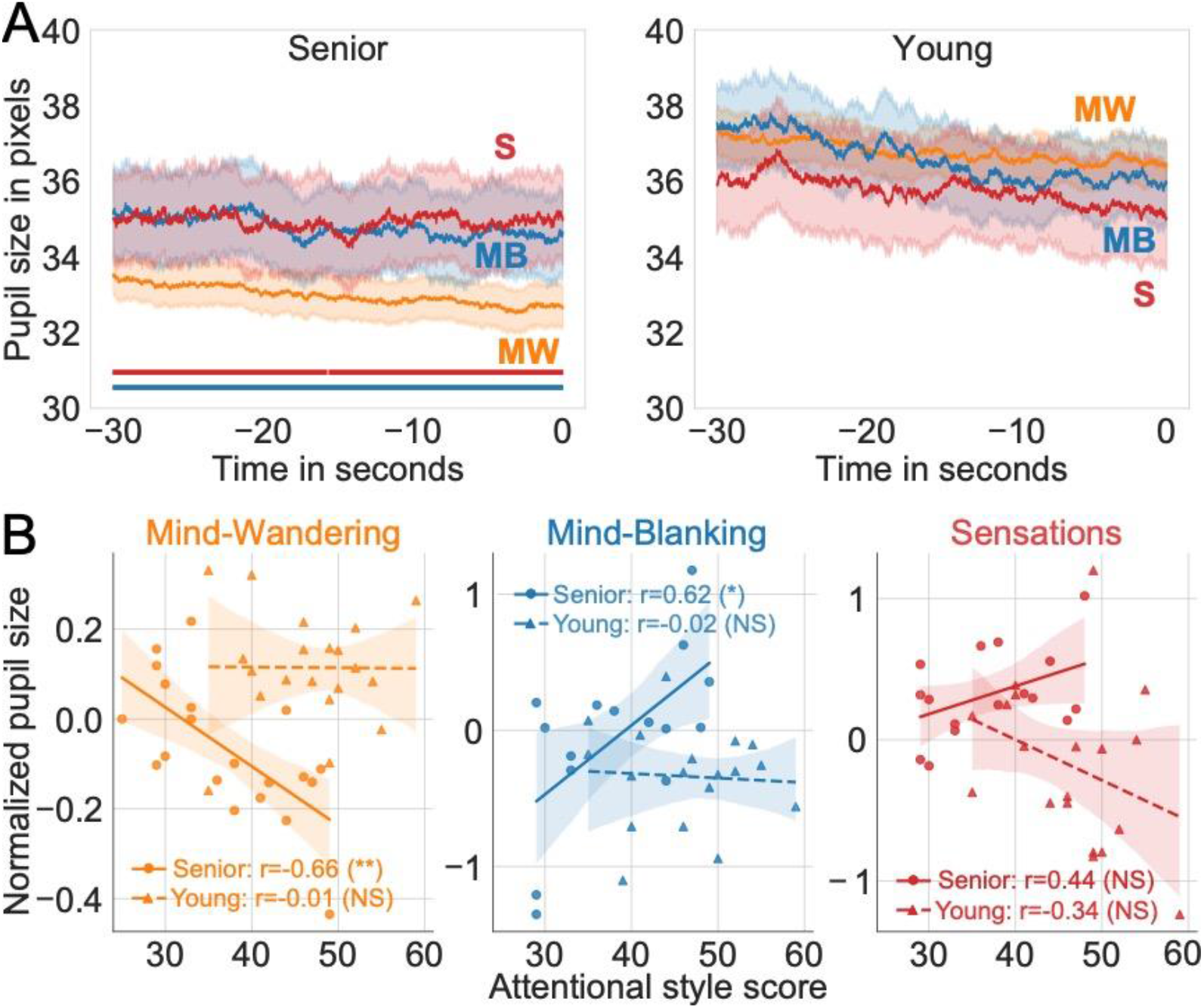
The relationship between pupil-linked arousal and ongoing mental state is dependent on attentional style in seniors but not in young participants. (A) The time-course of pupil size over thirty seconds preceding sound onset for each mental state showed that seniors had smaller pupil size when reporting mind wandering (MW) than mind blanking (MB) and sensory-related thoughts (S) (left panel). No such differences were found in young (right panel). *Notes*: Mean and confidence intervals are represented respectively with solid line and shaded areas. Clusters of significant difference between mental states are represented by solid lines (MW vs. MB: blue and MW vs. S: red). **(B)** Attentional style also interacted with age groups in predicting the pupil size differences, with less focused seniors showing lower pupil size for MW (left panel) and higher pupil size for MB (middle panel), while no such interaction was found in young. No significant correlations were found for S (right panel). *Notes*: Lines (straight: senior, dotted: young) represent the linear fit for each mental state and shaded areas are confidence intervals. Individual data points are represented as circles (senior) and triangles (young).

Attentional style furthermore interacted with age groups in predicting the pupil size difference between MW and MB (t(108,40)=-2.25, p=0.028) and between MW and S (t(108,40)=-2.33, p=0.023, Figure 4). Post-hoc analyses revealed that MW had overall larger normalized pupil size than MB in younger adults (t(17)=4.74, p<0.001, corrected for multiple comparison), and S than MW in seniors (t(15)=3.81, p=0.005). Importantly, attentional style correlated negatively with normalized pupil size in senior participants (Pearson’s correlation, R=-0.66, p=0.002), but not in young (R(17)=-0.01, p=0.971) (Figure 4, left panel). For MB, attentional style correlated positively with pupil size in senior participants (R(13)=0.62, p=0.015), but not in younger adults (R(16)=-0.063, p=0.803) (Figure 4, middle panel). Finally, a trend for a positive correlation between ASQ and pupil size was also found for S (R(14)=0.44, p=0.092, young: R(17)=-0.34, p=0.161) (Figure 4, right panel). Overall, these results indicate that the relationship between arousal and attentional style across ongoing mental states is specific to seniors, with the possibility that MB resulting from a hyper-arousal state which is translated as less focus.

Finally, we checked whether age-dependent changes in pupil size could be confounded by the variations of eyelid gap and gaze position that have been reported to also predict the occurrence of ongoing mental states (Grandchamp et al., 2014). Time-resolved analyses of eyelid gap and linear mixed models showed that MB was associated with a smaller eyelid gap in senior in comparison to S (t(40,108)=-2.32, p=0.236) and to MW (t(40,108)=-3.01, p=0.004) (Figure S5A). These differences interacted with ASQ scores but not with age (ASQ score for MB vs. MW: t(40,108)=3.47, p<0.001, ASQ score for MB vs. S: t(40,108)=2.61, p=0.011, Figure S5B). Gaze position did not predict mental states, nor depended on age and attentional style (Figure S6). These results show that other ocular metrics can add information regarding the occurrence of ongoing mental states and their interaction with attentional style, but not with age. Thus, they cannot confound the observed pupil size differences across mental states and the reported interaction between age and attentional style.

## Discussion

We investigated how age-dependent changes in attentional control and arousal impacts the regulation of ongoing mental states. Using an experience-sampling task at rest, overall we found that attentional control and arousal jointly regulate ongoing mental states in an age-dependent manner and uncover the presence a specific profile of ongoing mental state regulation in healthy aging. Our findings support the multi-faceted view of ongoing thinking, highlighting the need to carefully consider the age-dependent changes in the regulation of mental states when studying their psychological and physiological correlates (Robison et al., 2020).

We first show that both age groups had higher rates of Mind-Wandering (MW) over Mind-Blanking (MB) and Sensations (S), replicating previous findings. Indeed, such high prevalence of MW is consistent with evidence about MW occupying 10-60% of our mental activity during waking life (Killingsworth & Gilbert, 2010; Seli et al., 2018). We also replicated that MB is a quite rare phenomenon, reported between 8-18% depending on the thought sampling methodology (Ward & Wegner, 2013; Andrillon et al., 2021; Van Calster et al., 2017; Mortaheb et al., 2022). The low reports of Sensations can be explained by the limited visual and auditory stimulations (dim light, black screen and earphones) and task requirements (ongoing mental state reports at rest). The stable inter-state dynamics, distribution of mental states and reaction times across age (supplementary material) further indicate that ongoing mental states are discriminated similarly in both young and seniors. This pattern of results confirms the validity of our procedure in probing the dynamics of ongoing mental states across age groups at rest.

At the same time, seniors were overall more focused than younger participants, and showed an association between mental state reports and attentional control (Kraft et al., 2020; Van Calster et al., 2018), which was reversed to what we observed in young participants. Our results are in line with previous findings that seniors had better attentional control than younger adults (Maillet et al., 2020; Moran et al., 2021; Seli et al., 2021). The reversed relationships between attentional style and mental state rates indicate an expected association between lower attentional control and higher rates of MW in younger population, as young participants appear restless in general (Moran et al., 2021). Contrary to our prediction, though, lower attentional control was associated with lower rates of MW in seniors. These findings extend previous results showing strategic differences in how attentional control regulate the propensity to enter MW in seniors and younger adults (Moran et al., 2021). We finally replicated the findings that seniors exhibited overall lower anxiety symptoms than younger participants (Wolitzky-Taylor et al., 2010), and that it correlated with attentional style. However, we found no impact of this variable on ongoing mental states rates, showing importantly that trait-anxiety is not a confound for our reported effects.

By investigating pupil size, a proxy of arousal (Bradley et al., 2008; Unsworth & Robison, 2018), we further found that physiological signatures of ongoing mental states depend on attentional control in seniors, but not in young participants. In line with the literature, we observed that MW tended to be associated with higher arousal (pupil dilation), and MB with lower arousal (pupil restriction) in younger participants (Andrillon et al., 2021; Pelagatti et al., 2018; Unsworth & Robison, 2018). Conversely in seniors, attentional style modulated pupil size variations across mental states: higher arousal (pupil dilation) was associated with MW in focused seniors, and with MB in more distracted seniors. Such behavioral profile associating distractibility, higher arousal and higher prevalence of MB in seniors can also be found in other cases, for example under stress or conditions such as attention deficit hyperactivity disorder (Van den Driessche et al., 2017).

To further explain these latter effects, one might speculate on the neural origin of our findings, as pupil size variations are only an indirect measure of arousal. The age-dependent regulation of ongoing mental states by attentional control and arousal might be related to age-related changes in the activity of locus coeruleus (LC) during healthy aging (Jepma & Nieuwenhuis, 2011; Larsen & Waters, 2018). LC is a key brain region involved in the regulation of arousal (Sara & Bouret, 2012) and attentional control (Unsworth & Robison, 2017) and adjusts pupil size variations by subcortical pathways innervating the iris dilator and sphincter muscles (Mathôt, 2018). LC can also be involved in the regulation of mind wandering states as indicated by a neural model (Mittner et al., 2016). Thus, the interaction between pupil size and attentional style in seniors might be linked to the reduced connectivity of the LC observed during healthy aging (Langley et al., 2021). As structural and functional changes in LC (Chen et al., 2022), as well as impairment of attentional control (Baddeley, 2001), are also found in neurological conditions such as Alzheimer’s disease. Therefore, the probing of behavioral and (neuro)physiological correlates of spontaneous cognition might be useful to identify early markers of age-related neurological conditions (Kvavilashvili et al., 2020). Further studies using neuroimaging and active tasks, such as the auto-scrolling text (Ward & Wegner, 2013), are expected to identify the neural underpinnings and the behavioral relevance of age-dependent changes in the regulation of ongoing mental states. Investigating these mechanisms in age-related neurological conditions might constitute a promising route for identifying early prognostic markers of neurodegenerative diseases.

To conclude, we provide a comprehensive approach to ongoing mental states, including the newly introduced state of mind blanking. We uncover that ongoing mental states are regulated by attentional control and arousal differed in healthy seniors and younger adults.

## Author notes

### Contributions statement

A. de Beauchamp, C. Bastin and A. Demertzi designed the study. A. de Beauchamp and M. Koroma conducted the research and the analysis helped by S. Mortaheb and P.A. Boulakis. M. Koroma, A. de Beauchamp and A. Demertzi wrote the manuscript. All authors reviewed the manuscript.

### Data and code availability

All codes and data are freely accessible via the Gitlab (https://gitlab.uliege.be/S.Mortaheb/mb_aging) and OSF server (https://osf.io/mknjr/).

