## Supplementary Information for "Elsewhere or Blanked? Ongoing mental states are regulated by pupil-linked arousal and attentional style in healthy ageing"

**Short title**: Age-dependent regulation of spontaneous thinking

Matthieu Koroma[1,2], Aurèle Robert de Beauchamp[1], Sepehr Mortaheb[1,2], Paradeisios Alexandros Boulakis[1,2], Christine Bastin[1,2,3] and Athena Demertzi[1,2,3*]

[1] GIGA-CRC In Vivo Imaging, Allée du 6 Août, 8 (B30), 4000 Sart Tilman, University of Liège BELGIUM
[2] Fund for Scientific Research FNRS, Brussels, Belgium
[3] Department of Psychology, , University of Liège, Liège, Belgium

**Keywords**: Attention, Mind-blanking, Mind-wandering, Pupillometry, Arousal, Aging

***Corresponding author**:
Physiology of Cognition
GIGA-CRC In Vivo Imaging Center
Allée du 6 Août, 8 (B30), 4000 Sart Tilman
University of Liège
BELGIUM


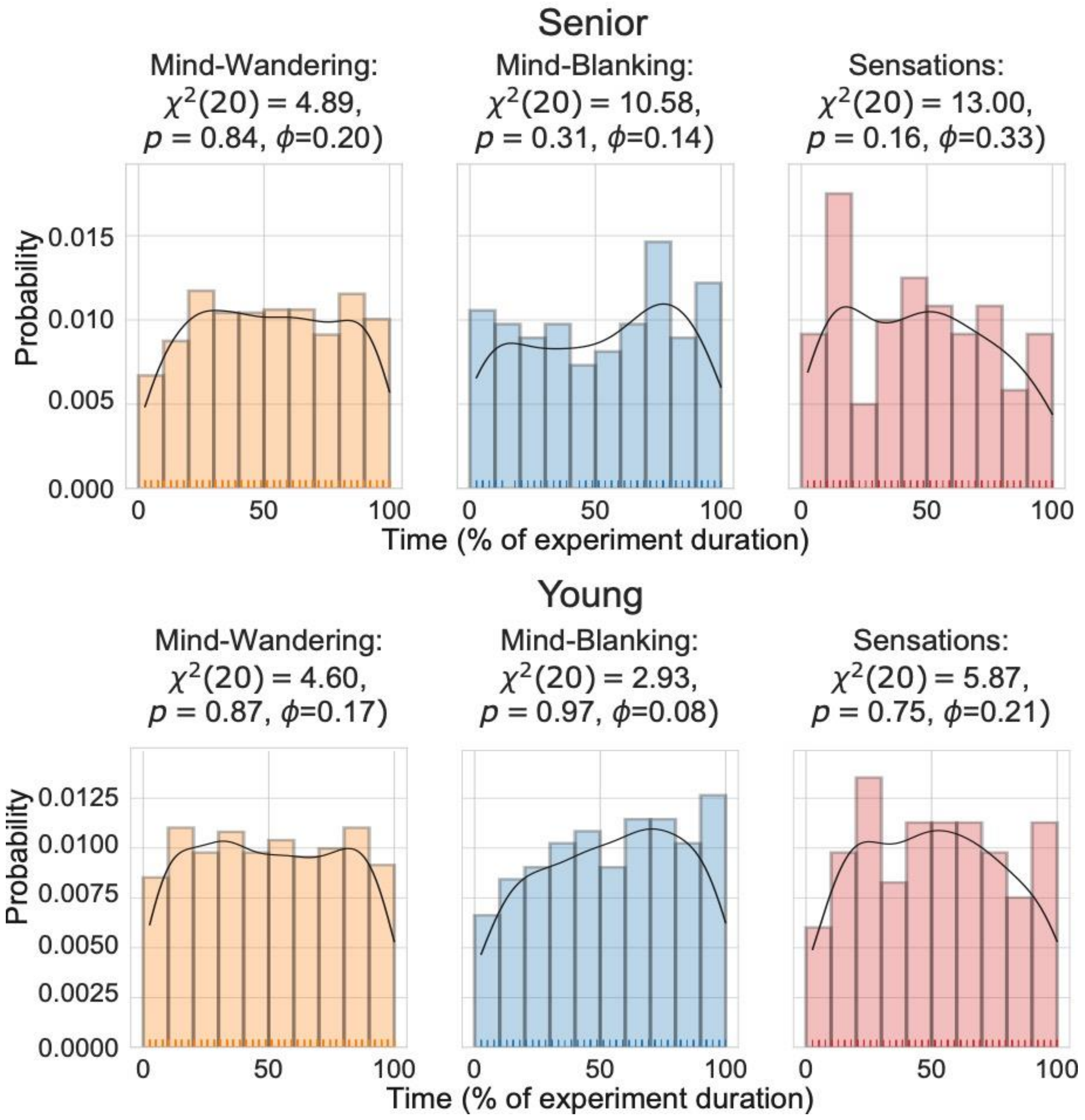


**Figure S1. Probability of responses does not vary with the duration of the experiment**. The histograms composed of 10 bins related to the equal divisions of experiment duration to its 10% time slots for each mental states. Notes: left panel, yellow: Mind-Wandering; middle panel, blue: Mind-Blanking, right panel, red: Sensations, in senior (upper panels) and young (lower panels) participants. Solid line represents the fit across the whole experiment of the probability of response, plotted for each condition. Testing for the null hypothesis of uniform distribution of probability of mental state (Chi-squared test) is reported at the top of each panel.

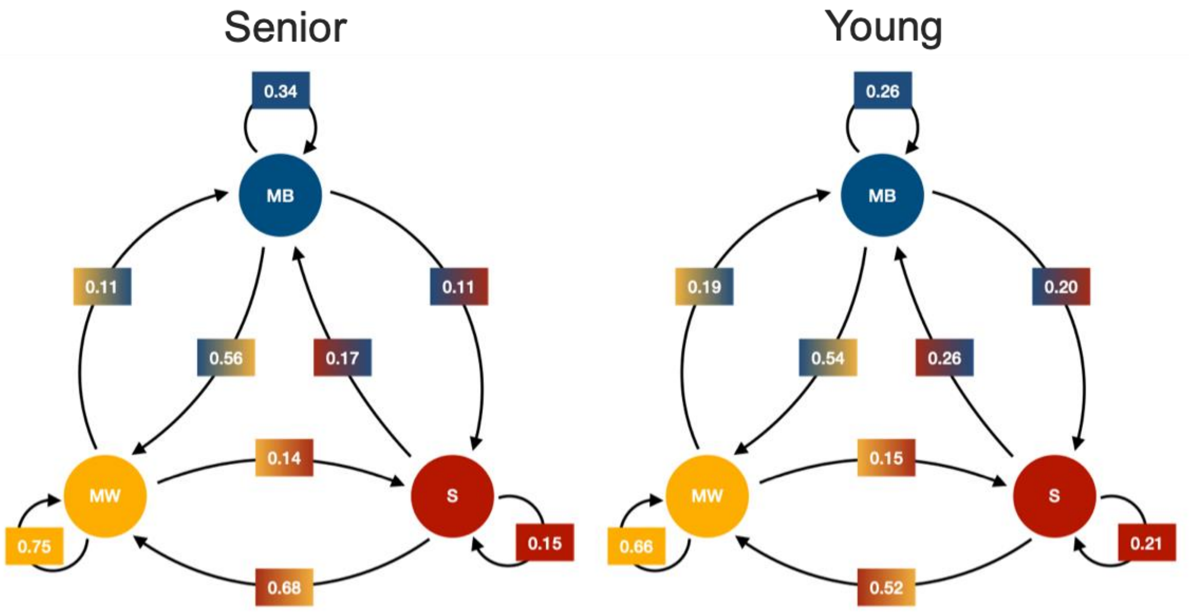


**Figure S2. Inter-state dynamics across mental states do not vary across age groups.** The probability of transitions across mental states were similar across participants, with no evidence for a significant different across age groups (left panel: senior, right panel: young) (Wilcoxon rank-sum test, $p>0.05$ for all tests even before multiple comparisons correction). Notes: Mind-Wandering (MW): yellow, Mind-Blanking (MB): blue, Sensations (S): red.

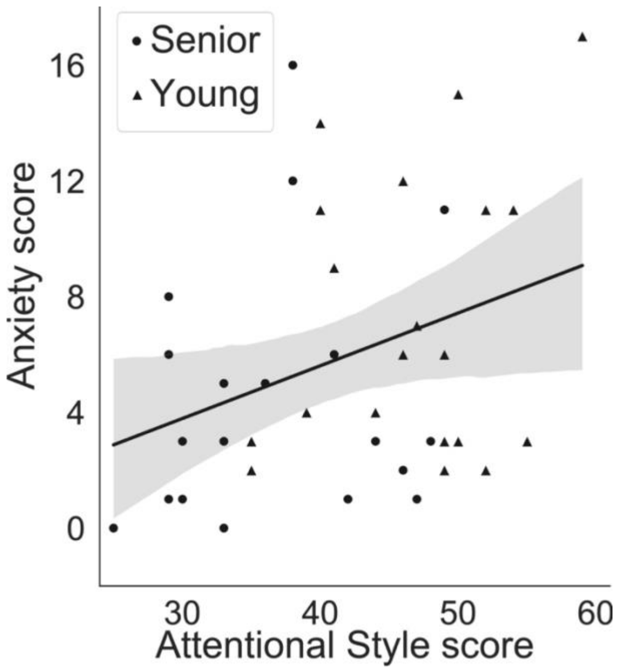


**Figure S3. Anxiety and attentional style scores correlate positively, in that less attentional focus is linked to higher anxiety scores**. The line represents the linear fit between the Attentional Style Questionnaire (ASQ) score and Generalized Anxiety Disorder (GAD) with shaded areas as confidence intervals. Individual data points are represented as circles (senior) and triangles (young).

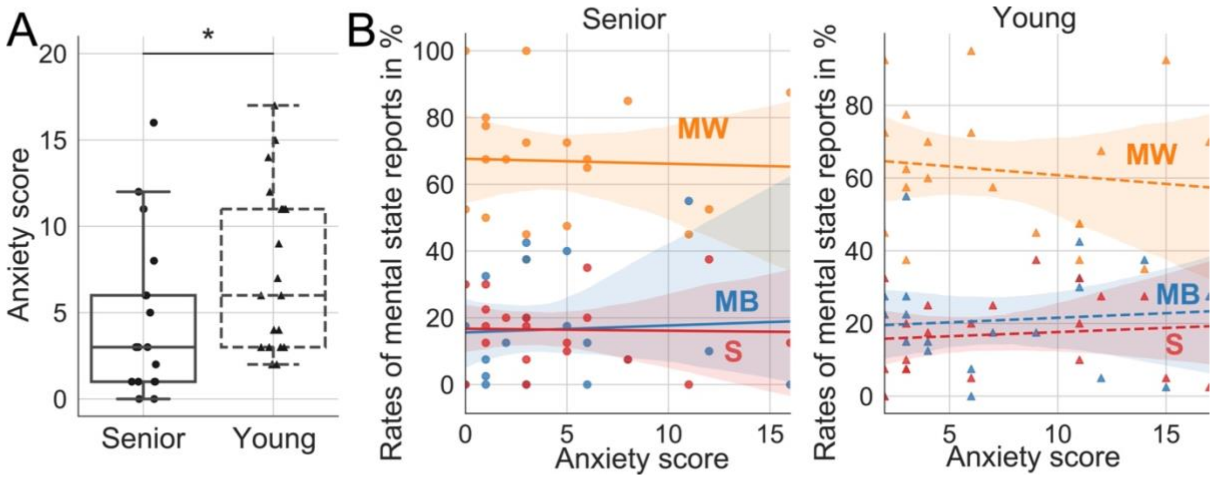


**Figure S4. Age-dependent anxiety does not impact the reportability of spontaneous mental states. (A)** Seniors self-reported lower scores on the Generalized Anxiety Disorder (GAD) than younger participants, indicative of lower anxiety symptoms. (**B**) No association between GAD and attentional style nor interaction with age in predicting mental state reportability were found, excluding anxiety as a potential confound explaining the influence of attentional style on mental state reportability across age groups. Notes: Boxplots: mean (horizontal line) and first and third quartile (box), as well as first and ninth deciles (vertical line) of the GAD score are represented (solid: aged, dotted: young). Individual data points are represented in circles: (senior) and triangles (young). Significant differences across conditions are reported as solid black lines (Wilcoxon rank-sum test, ***p<0.001). Correlation plots: The lines (solid: senior, young: dotted) represent the correlation between the rate of Mind-Wandering (MW): yellow, Mind-Blanking (MB): blue, Sensations (S): red, and the GAD score with shaded area as confidence intervals for both senior (left panel) and young (right panel) participants. Individual data points of each are represented as circles (senior) and triangles (young).

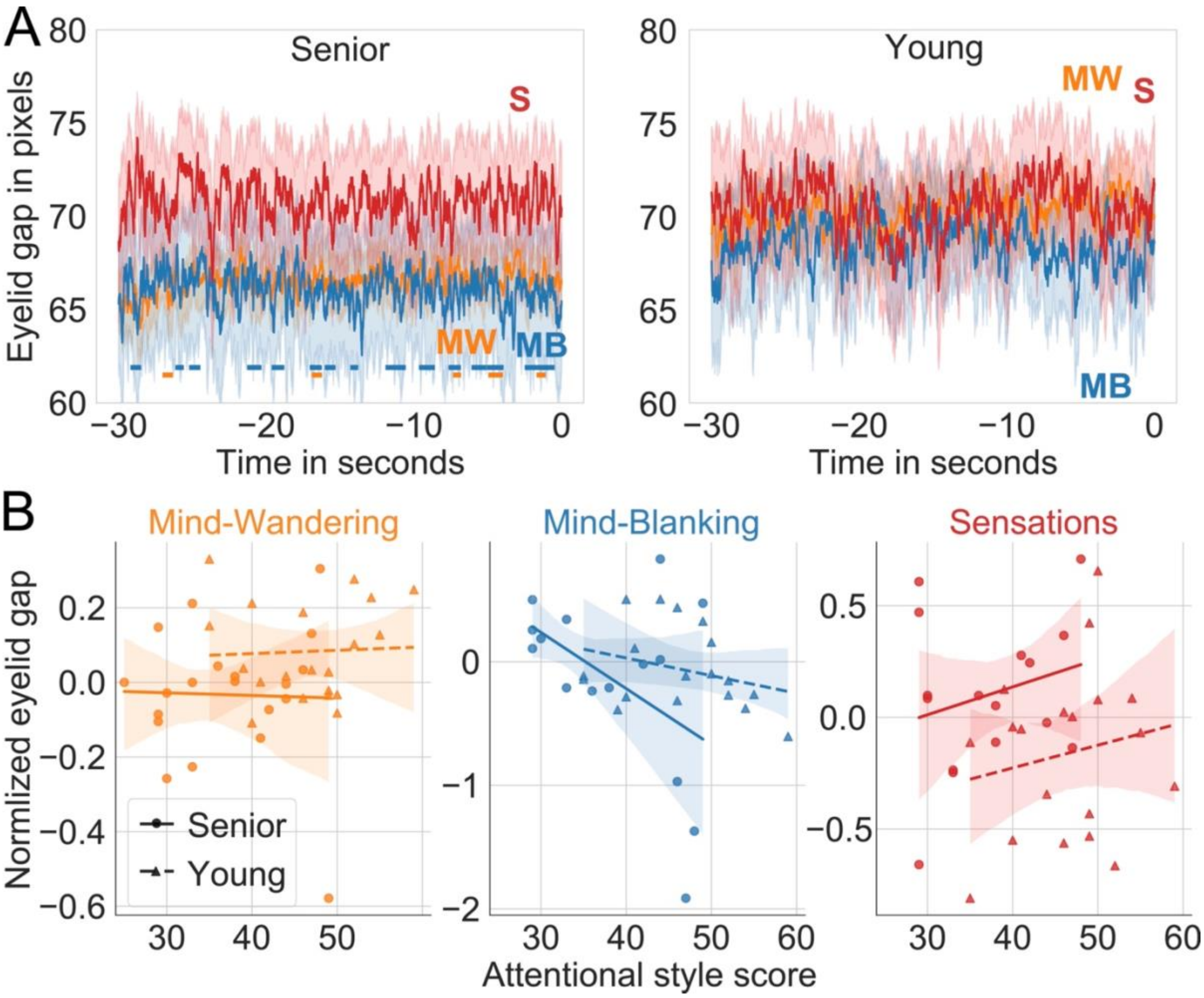


**Figure S5**. **Eyelid gap variations across spontaneous mental states do not interact with attentional style and age. (A)** The time-course of eyelid gap over thirty seconds preceding sound onset for each mental state showed that seniors had smaller eyelid gap size when reporting mind wandering (MW) and mind blanking (MB) compared to sensory-related thoughts (S) (left panel). No such differences were found for young (right panel). Notes: Mean and confidence intervals are represented respectively with solid line and shaded areas. Clusters of significant difference between mental states are represented by solid lines (MW vs. S: yellow and MB vs. S: blue). **(C)** Attentional style did not interact with age groups in predicting eyelid gap variations. Notes: Lines (straight: senior, dotted: young) represent the linear fit for each mental state and shaded areas are confidence intervals. Individual data points are represented as circles (senior) and triangles (young).

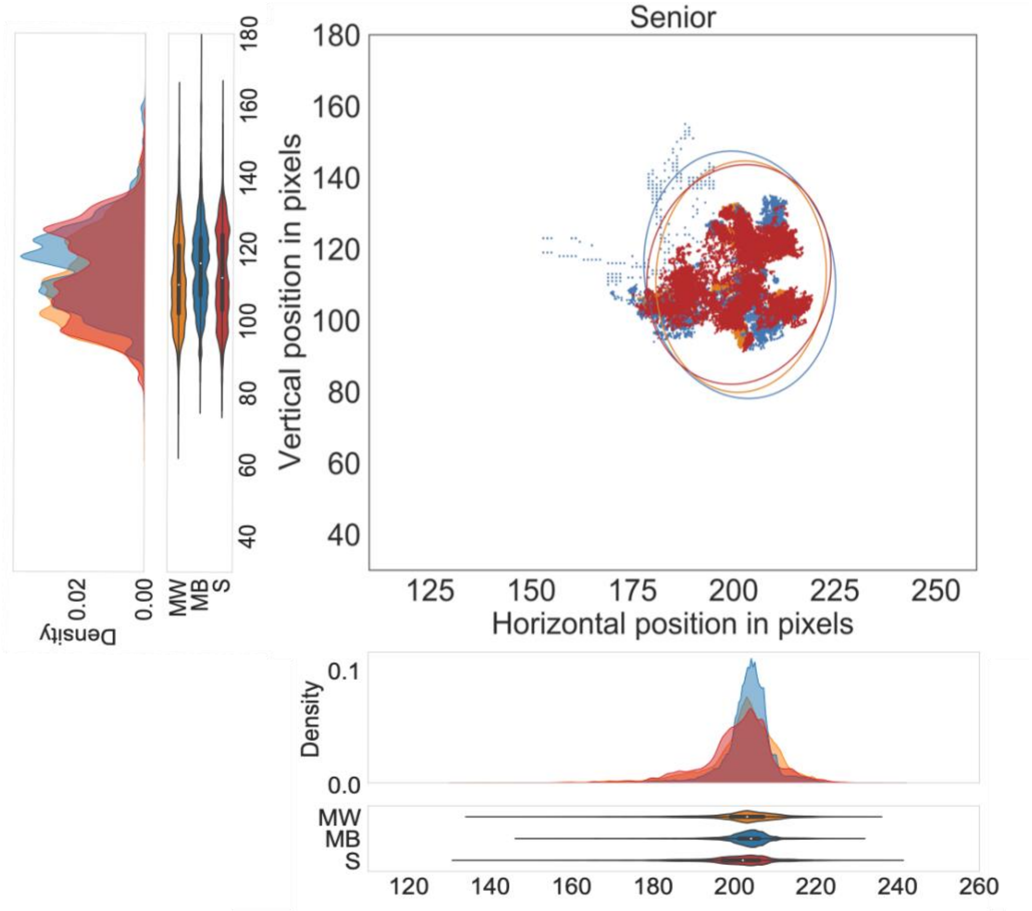
Senior
Vertical position in pixels
Horizontal position in pixels
Density
MW
MB
S

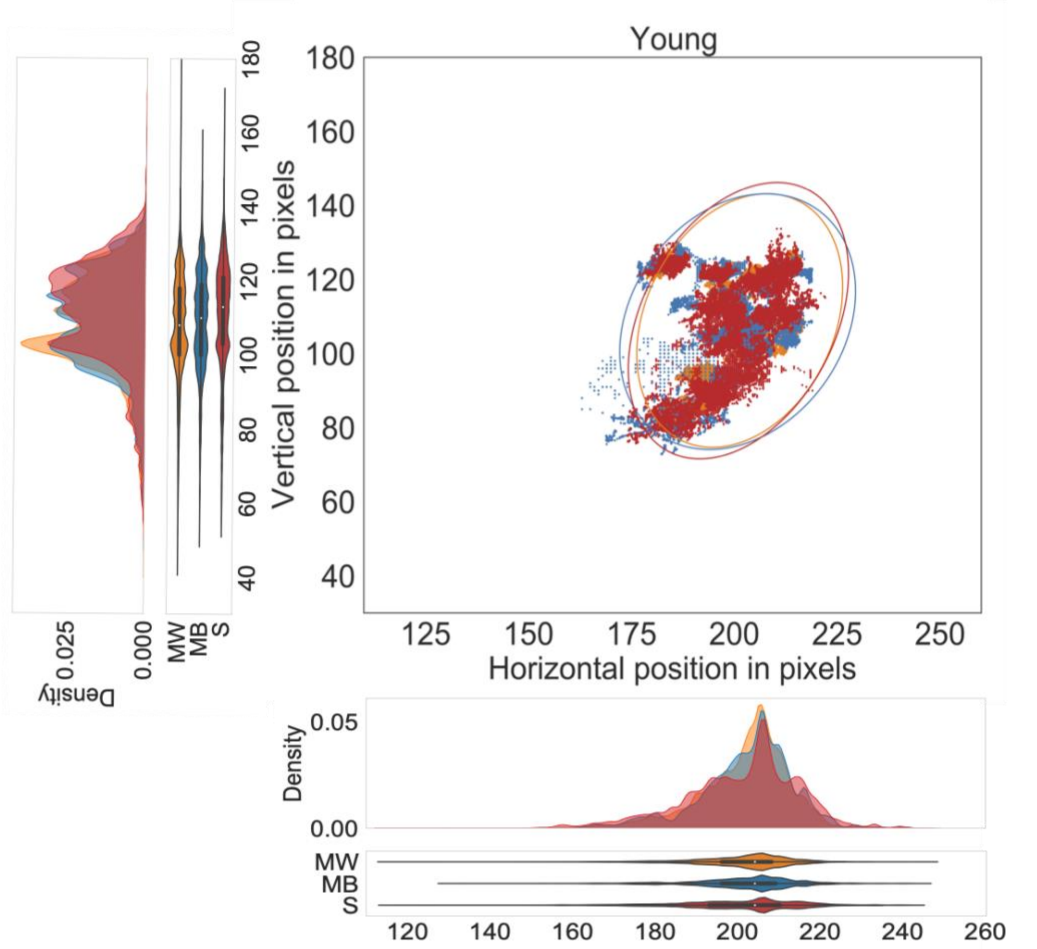
Young
Vertical position in pixels
Horizontal position in pixels
Density
MW
MB
S

**Figure S6. Gaze position is not modulated by mental state neither in senior nor young participants.** The position of the gaze over thirty seconds preceding sound onset for Mind-Wandering (MW): yellow, Mind-Blanking (MB): blue, Sensations (S): red, in senior (upper panels) and young (lower panels) participants are represented. Notes: mean and confidence intervals are represented respectively with solid line and ellipse. Distribution and violin plots of the horizontal (bottom panels) and vertical (left panels) gaze position over the thirty seconds preceding sound probe for each mental state.
